# Feed restriction in mid-lactation dairy cows. II: Effects on protein metabolism-related blood metabolites

**DOI:** 10.1101/2020.06.12.144980

**Authors:** I. Ansia, Y. Ohta, T. Fujieda, J. K. Drackley

## Abstract

The aim of the study was to describe the metabolic responses of protein metabolism to a period of negative nutrient balance induced by feed restriction (FR). Seven multiparous Holstein cows (93 ± 15 days in milk) were randomly assigned to 7 treatments in a 7 × 4 Youden square design. Daily intake was restricted to provide 60% of energy requirements during 5 d except for one treatment with ad libitum (AL) feeding. While 5 out of 7 experimental treatments involved abomasal supplementation of AA or glucose, in this article we evaluated only the effects of FR by comparing both control treatments (AL and FR). Data of 2 cows within the AL group were removed due to sickness and therefore it had n = 2. A rapid decrease of most amino acids in plasma was paired with an increase in blood urea N with its peak on d 2 and decreasing afterwards. On the other hand, Lys, Arg, Gly, Gln, and Cys were greater during FR. Comparing the fluctuation of all the essayed N components in circulation across the 5-d period, protein tissue mobilization may have supplied amino acids for catabolism to provide needs for N and energy precursors.

**Implications:** The short-term feed restriction model described in this article can serve as an alternative to study metabolic adaptations during the transition period. The response observed of the protein metabolism sets the baseline to measure the effect of nutrients supplementation and identify those candidates that will improve milk production and overall health after calving.

## Introduction

Distribution and efficient use of the required amino acids (**AA**) during a period of lower DMI requires an inter-organ coordination between liver and peripheral tissues (Arriola Apelo et al., 2014). Amino acid uptake by the mammary gland assumes a high priority because of the relative amount consumed in comparison with the splanchnic tissues (Hanigan et al., 2001), and because the mammary gland is able to alter extraction efficiency according to its requirements (Mepham, 1982; Mackle et al., 2000). Accordingly, the negative protein balance during the transition period is reflected in lower concentrations of the majority of essential and non-essential AA in circulation, which eventually will recover their “pre-transition” values later during lactation (Maeda et al., 2012). Nevertheless, analysis of the AA individually during the transition period can sometimes lead to misleading conclusions due to the effect of different feeding or grouping strategies applied (Doepel et al., 2002).

The objective of this study was to characterize and quantify the effects of feed restriction on energy (Ansia et al., unpublished results) and protein metabolism. Our hypothesis was that timing and magnitude of the variation in certain blood metabolites could highlight the most important metabolic pathways and accentuate requirements for specific AA in maintenance of milk production and basal metabolism when DMI is deficient. In a companion paper we reported that a feed restriction to 60% of the net energy requirements for lactation decreased milk production, and milk protein and lactose yield. Feed restriction also elicited a metabolic response characterized by an increase of NEFA paired with a decrease in glucose and insulin in circulation. In addition, feed restriction triggered a decrease in albumin and an increase in bilirubin and triglycerides concentrations (Ansia et al., unpublished results). Our specific objective of this analysis is to set the baseline to evaluate the effects of specific nutrient supplementation within this model.

## Materials and methods

### Animals and diets

A more detailed description of the materials and methods used in this experiment can be found in a companion paper (Ansia et al., unpublished results). Briefly, seven rumen-cannulated multiparous Holstein cows past peak lactation (93 ± 15 d in milk; BW = 674 ± 43 kg) were used in a 7 × 4 Youden square design with 4 experimental periods of 10 d. Seven treatments were applied to evaluate the effects of 5 different abomasal infusions of macronutrients on energy and protein metabolism during a 5-d feed restriction period. Only the results from the comparison of the 2 treatments used as a positive (ad libitum; **AL**) and negative (feed restriction; **FR**) controls are discussed in this article and will be referred to as diets. Cows during AL were fed to ensure daily orts of >15 % of the feed amount offered, and FR cows were fed to meet 60% of their net energy requirements for lactation (NE_L_) at the beginning of each period. During the following 5 d (d 6 to 10) all 7 cows were fed for ad libitum DMI to obtain 15% of refusals as fed, as a recovery and wash-out period. The diet, fed as a total mixed ration, was described elsewhere (Ansia et al., unpublished results).

### Blood Sampling and Analysis

Indwelling catheters were inserted into a jugular vein of each cow on the day before d 1 of each period. During d 1 to d 5, blood samples were obtained at -1, 0, 3, 4, 5 and 14 h relative to feeding each day. Catheters were flushed with sterile heparinized saline every 6 h between daily sampling periods and were removed on d 5 in each period. Protein metabolism was assessed by analyzing concentrations of all EAA and non-essential AA (NEAA), blood urea N (BUN), and other metabolites of the N cycle. Concentrations in deproteinized plasma were measured by Ajinomoto Co., Inc. (Tokyo, Japan) using an L-8900 AA analyzer (Hitachi, Tokyo, Japan). The incremental area under the curve (IAUC) of the daily average across the 5-d period was calculated with the trapezoid method for each AA and BUN.

### Statistical Analysis

Two cows were diagnosed with a bacterial endotoxemia and clinical mastitis, respectively, both within the AL group. Due to the noticeable detrimental effect observed on milk production, DMI, and other blood variables, and the influence that their data had on the normality and homogeneity of the model, we decided to remove them from all statistical analysis of performance and blood constituents.

Comparisons of all variables between treatments were made using the MIXED procedure of SAS version 9.4 (SAS Institute, 2012). Effects of treatment, period, day, and hour were included as fixed effects while cow and the interaction cow by treatment were included as a random effects. The repeated fixed effect of day nested within period was used in the REPEATED statement with cow nested in treatment as the subject. The CONTRAST statement was used to compare AL and FR. The linear and quadratic trends were evaluated using the ESTIMATE statement from PROC MIXED, to classify the evolution of the daily average across the period.

The PROC UNIVARIATE and the INFLUENCE option within PROC MIXED were applied to each variable to check for normality of residuals and for the presence of outliers. When appropriate, variables were transformed to accomplish the above-mentioned criteria. The proper power transformation was selected according with the convenient lambda displayed by the TRANSREG procedure. Non-transformed LSMEANS and SEM were reported.

As a complementary analysis to simplify detection of interrelationships among N compounds involved in protein metabolism, we performed factor analysis using PROC FACTOR in SAS after standardization of the data set. We used the principal component method to estimate the number of parameters, and the varimax rotation for a clearer interpretation of the factor loadings. The final number of factors is selected by dimensionality assessment according with the Kaiser’s criterion eigenvalue > 1.00 and graphically according with the scree plot generated by this procedure. A factor loading was considered for discussion only when higher than a cut-off value of 0.5. When a variable belonged to more than one factor, it was included in the one where it had the greater loading.

## Results

The FR diet reduced daily plasma concentrations of Asp (*P*_Diet_ < 0.01), Tyr (*P*_diet_ < 0.01), Pro (*P*_Diet_ = 0.01), and tended to reduce it for Thr (*P*_Diet_ = 0.08), His (*P*_Diet_ = 0.06) and Glu (*P*_Diet_ = 0.07; Table 1). Despite a lack of diet effect, concentrations of other AA such as Ala, Ser, Ile, Val, Trp, Asn, and Met also decreased (*P*_FR x Day_ ≤ 0.01) across days only during FR (Table 2). All these AA showed or tended to show a quadratic decline, whereas for Asp, Ile and Met the decrease was linear. Contrarily, plasma concentration was higher during FR for Arg (*P*_Diet_ = 0.05) and tended to be higher for Gly (*P*_Diet_ = 0.09), and Lys (*P*_Diet_ = 0.10). Concentrations of individual AA were lower during the postprandial sampling (data not shown). Only the NEAA showed a significant decrease (Figure 1). There were no differences among diets except for Glu, which was higher after feeding (data not shown).

**Table 1.**
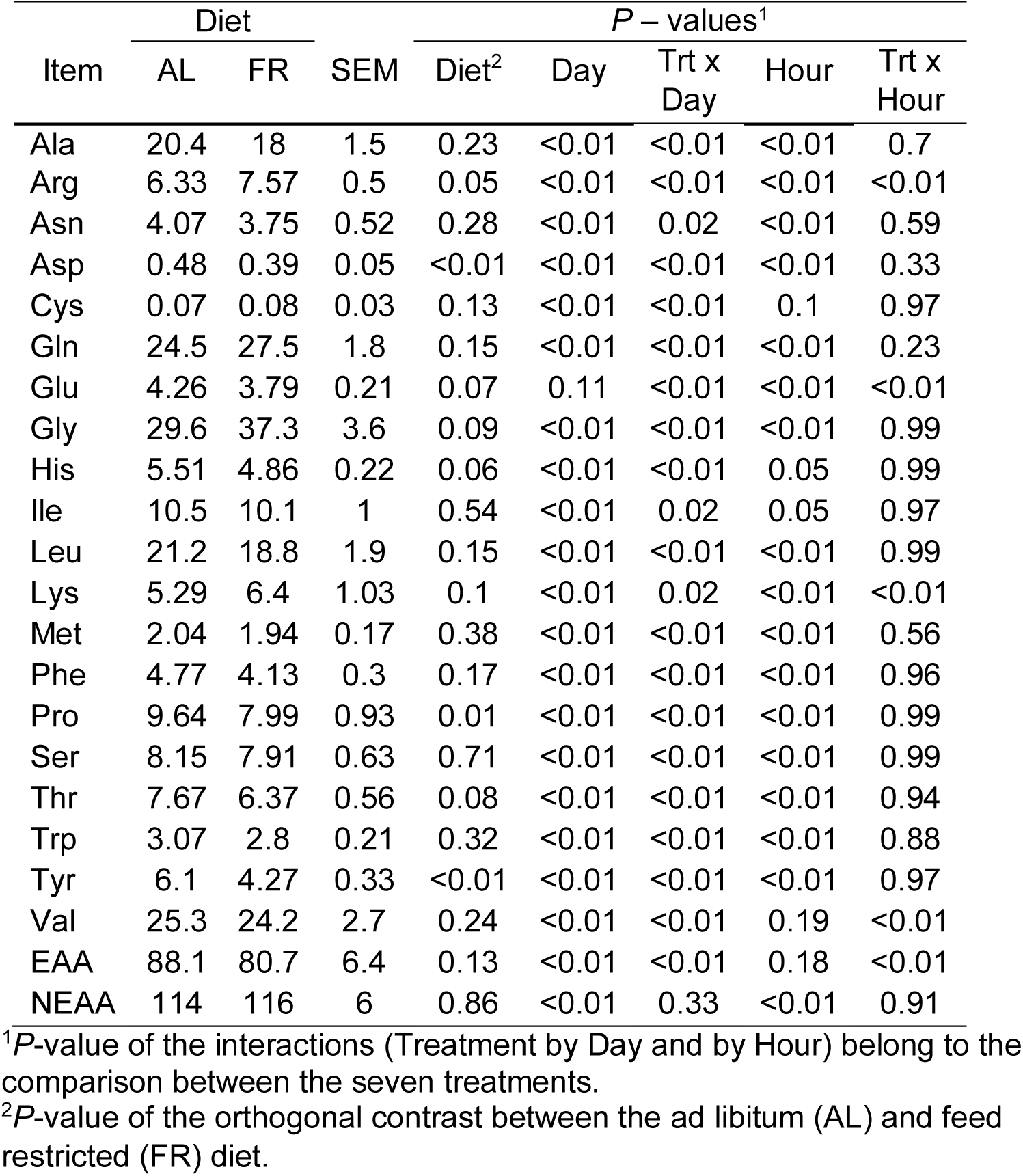
Least squares means of the concentrations (μmol/dL) of blood amino acids of mid-lactation dairy cows during 5 days on the ad libitum (AL) and feed-restricted (FR) diets.

**Table 2.** Least squares means of the concentrations (μmol/dL) of blood amino acids of mid-lactation dairy cows during 5 days on the ad libitum (AL) and feed-restricted (FR) diets per day.

| Item | Diet | Day |  |  |  |  | SEM | P-values |  |  |
| --- | --- | --- | --- | --- | --- | --- | --- | --- | --- | --- |
|  |  | 1 | 2 | 3 | 4 | 5 |  | Diet x Day<sup>3</sup> | Trends<sup>2</sup> |  |
|  |  |  |  |  |  |  |  |  | Lin. | Quad |
| Ala | AL | 22.5 | 21.1 | 20.9 | 18.9 | 18.5 | 1.74 | 0.41 | 0.02 | 0.94 |
|  | FR | 21.6<sup>a</sup> | 16.8<sup>b</sup> | 16.8<sup>b</sup> | 17.7<sup>b</sup> | 17.1<sup>b</sup> | 1.19 | <0.01 | 0.01 | <0.01 |
| Arg | AL | 6.79 | 6.43 | 6.27 | 5.94 | 6.23 | 0.62 | 0.64 | 0.19 | 0.43 |
|  | FR | 7.43 | 7.49 | 7.49 | 7.93 | 7.53 | 0.48 | 0.68 | 0.45 | 0.64 |
| Asn | AL | 4.19 | 4.39 | 4.45 | 4.24 | 3.87 | 0.39 | 0.65 | 0.46 | 0.17 |
|  | FR | 5.06<sup>a</sup> | 3.68<sup>b</sup> | 3.41<sup>b</sup> | 3.58<sup>b</sup> | 3.15<sup>b</sup> | 0.33 | <0.01 | <0.01 | <0.01 |
| Asp | AL | 0.47 | 0.46 | 0.51 | 0.47 | 0.52 | 0.06 | 0.76 | 0.48 | 0.9 |
|  | FR | 0.51<sup>c</sup> | 0.44<sup>bc</sup> | 0.36<sup>ab</sup> | 0.31<sup>a</sup> | 0.31<sup>a</sup> | 0.05 | <0.01 | <0.01 | 0.41 |
| Cys | AL | 0.08<sup>a</sup> | 0.09<sup>a</sup> | 0.05<sup>b</sup> | 0.05<sup>c</sup> | 0.07<sup>b</sup> | 0.03 | <0.01 | 0.24 | 0.21 |
|  | FR | 0.08<sup>a</sup> | 0.08<sup>a</sup> | 0.07<sup>b</sup> | 0.08<sup>a</sup> | 0.09<sup>a</sup> | 0.02 | 0.01 | 0.44 | 0.15 |
| Gln | AL | 25 | 24.2 | 25.2 | 24.5 | 23.6 | 2.1 | 0.78 | 0.65 | 0.64 |
|  | FR | 27.4<sup>a</sup> | 25.7<sup>b</sup> | 27.6<sup>a</sup> | 28.6<sup>a</sup> | 28.1<sup>ab</sup> | 1.59 | 0.09 | 0.28 | 0.61 |
| Glu | AL | 4.2 | 4.38 | 4.18 | 4.09 | 4.47 | 0.26 | 0.37 | 0.67 | 0.45 |
|  | FR | 4.16<sup>a</sup> | 3.71<sup>b</sup> | 3.71<sup>b</sup> | 3.60<sup>b</sup> | 3.77<sup>b</sup> | 0.2 | 0.01 | 0.04 | 0.01 |
| Gly | AL | 30.9 | 29.6 | 29.5 | 29.8 | 27.9 | 3.98 | 0.81 | 0.54 | 0.91 |
|  | FR | 35.6<sup>a</sup> | 35.4<sup>b</sup> | 36.6<sup>b</sup> | 41.4<sup>b</sup> | 37.4<sup>b</sup> | 3 | <0.01 | 0.16 | 0.39 |
| His | AL | 5.67<sup>a</sup> | 5.90<sup>a</sup> | 5.80<sup>a</sup> | 5.46<sup>a</sup> | 4.74<sup>b</sup> | 0.34 | 0.01 | 0.03 | 0.01 |
|  | FR | 5.93<sup>a</sup> | 4.75<sup>b</sup> | 4.42<sup>c</sup> | 4.65<sup>bc</sup> | 4.52<sup>bc</sup> | 0.26 | <0.01 | <0.01 | <0.01 |
| Ile | AL | 11.7 | 11.1 | 10.8 | 9.5 | 9.3 | 1.1 | 0.3 | 0.03 | 0.91 |
|  | FR | 11.4<sup>a</sup> | 10.0<sup>b</sup> | 9.7<sup>b</sup> | 10.0<sup>b</sup> | 9.3<sup>b</sup> | 0.9 | 0.01 | 0.03 | 0.25 |
| Leu | AL | 22.8 | 22.5 | 22.2 | 19.9 | 18.5 | 2.1 | 0.21 | 0.03 | 0.31 |
|  | FR | 23.3<sup>a</sup> | 18.6<sup>b</sup> | 17.5<sup>b</sup> | 17.9<sup>b</sup> | 16.7<sup>b</sup> | 1.6 | <0.01 | <0.01 | <0.01 |
| Lys | AL | 6.16 | 5.41 | 5.19 | 4.79 | 5.01 | 1.28 | 0.55 | 0.44 | 0.58 |
|  | FR | 6.54 | 6.31 | 6.15 | 6.7 | 6.29 | 0.8 | 0.36 | 0.96 | 0.88 |
| Met | AL | 2.11 | 2.14 | 2.22 | 1.89 | 1.88 | 0.2 | 0.42 | 0.07 | 0.26 |
|  | FR | 2.23<sup>a</sup> | 1.89<sup>bc</sup> | 1.91<sup>bc</sup> | 1.92<sup>b</sup> | 1.78<sup>c</sup> | 0.14 | <0.01 | <0.01 | 0.2 |
| Phe | AL | 5.13 | 5 | 4.8 | 4.3 | 4.64 | 0.42 | 0.23 | 0.07 | 0.47 |
|  | FR | 5.14<sup>a</sup> | 3.95<sup>b</sup> | 3.70<sup>b</sup> | 3.92<sup>b</sup> | 3.95<sup>b</sup> | 0.33 | <0.01 | <0.01 | <0.01 |
| Pro | AL | 9.91 | 9.85 | 10.18 | 9.62 | 8.65 | 0.97 | 0.32 | 0.21 | 0.15 |
|  | FR | 9.78<sup>a</sup> | 7.73<sup>b</sup> | 7.41<sup>b</sup> | 7.88<sup>b</sup> | 7.17<sup>b</sup> | 0.73 | <0.01 | <0.01 | 0.01 |
| Ser | AL | 8.42 | 8.59 | 8.56 | 8.07 | 6.93 | 0.75 | 0.12 | 0.1 | 0.05 |
|  | FR | 9.06<sup>a</sup> | 7.65<sup>bc</sup> | 7.31<sup>c</sup> | 7.93<sup>b</sup> | 7.44<sup>bc</sup> | 0.56 | <0.01 | 0.1 | 0.05 |
| Thr | AL | 7.79 | 7.73 | 7.99 | 7.61 | 7.25 | 0.67 | 0.79 | 0.47 | 0.42 |
|  | FR | 7.85<sup>a</sup> | 5.82<sup>b</sup> | 5.83<sup>b</sup> | 6.35<sup>b</sup> | 6.00<sup>b</sup> | 0.53 | <0.01 | 0.01 | <0.01 |
| Trp | AL | 3.12 | 2.98 | 2.99 | 3.06 | 3.22 | 0.24 | 0.69 | 0.59 | 0.18 |
|  | FR | 3.21<sup>a</sup> | 2.83<sup>b</sup> | 2.63<sup>b</sup> | 2.73<sup>b</sup> | 2.63<sup>b</sup> | 0.18 | <0.01 | <0.01 | 0.01 |
| Tyr | AL | 6.4 | 6.45 | 6.39 | 5.76 | 5.53 | 0.45 | 0.69 | 0.11 | 0.38 |
|  | FR | 6.76<sup>a</sup> | 4.54<sup>b</sup> | 3.57<sup>c</sup> | 3.39<sup>cd</sup> | 3.08<sup>d</sup> | 0.32 | <0.01 | <0.01 | <0.01 |
| Val | AL | 29.5 | 28.7 | 28.9 | 26.6 | 24.7 | 3.4 | 0.31 | 0.14 | 0.4 |
|  | FR | 28.8<sup>a</sup> | 23.0<sup>b</sup> | 21.5<sup>b</sup> | 22.3<sup>b</sup> | 21.8<sup>b</sup> | 2.5 | <0.01 | 0.01 | <0.01 |
| NEAA | AL | 120 | 116 | 117 | 112 | 107 | 8 | 0.54 | 0.16 | 0.62 |
|  | FR | 125<sup>a</sup> | 110<sup>b</sup> | 112<sup>ab</sup> | 119<sup>ab</sup> | 112<sup>ab</sup> | 6 | <0.01 | 0.26 | 0.09 |
| EAA | AL | 93.8 | 91.5 | 91.5 | 83.7 | 80 | 7.4 | 0.34 | 0.05 | 0.51 |
|  | FR | 96.1<sup>a</sup> | 78.9<sup>b</sup> | 75.1<sup>b</sup> | 78.4<sup>b</sup> | 75.2<sup>b</sup> | 6.2 | <0.01 | <0.01 | <0.01 |

<sup>1</sup>P-value of the interaction of each diet (AL and FR) with day. Extracted from the SLICE statement output within the interaction of treatment with day.
<sup>2</sup> P-values of the test for linear or quadratic trends.
Means in a row with superscripts without a common letter differ, $P \leq 0.05$ .

**Figure 1.**
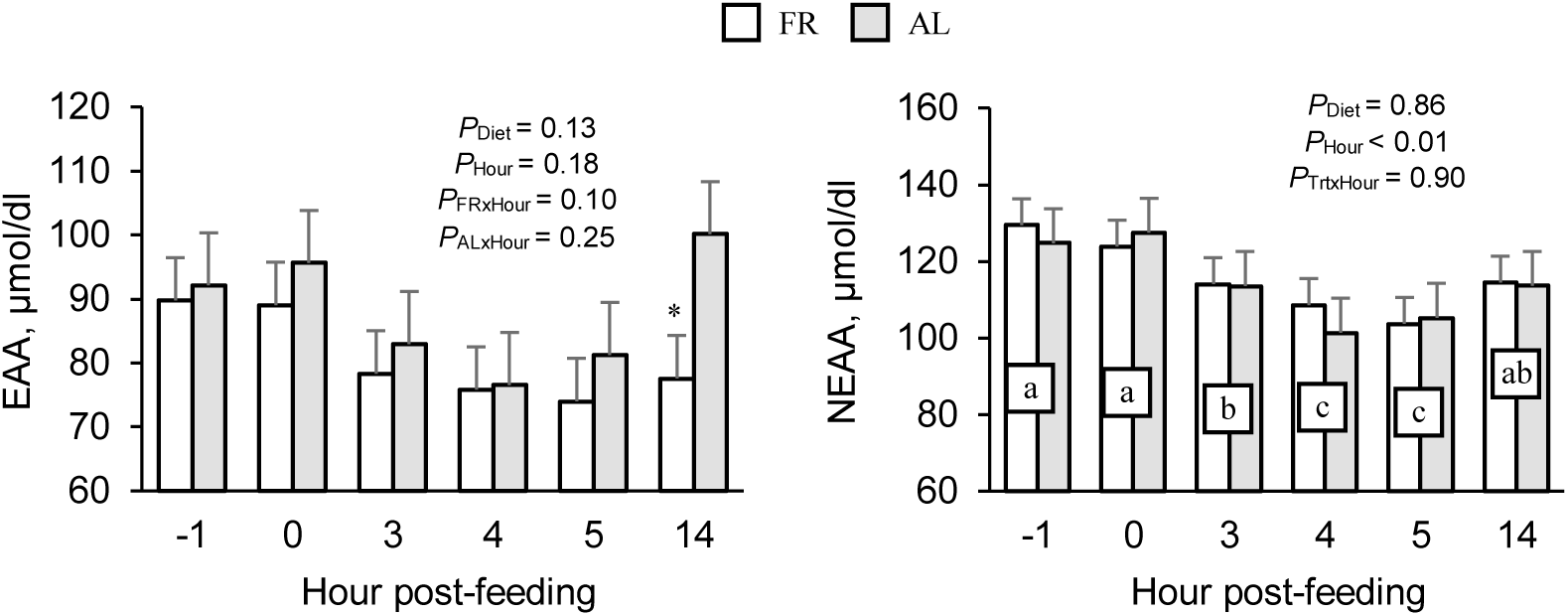
Concentrations of EAA and NEAA in plasma averaged across the 5-d period at -1, 0, +3, +4, +5 and +14 h after feeding during ad libitum (AL) and feed restricted diets (FR). Concentrations at same time points with different letter are significantly different. Differences between diets are marked at each time point with † (*P* < 0.10) and * (*P* < 0.05).

Ammonia (NH_3_) concentration in plasma was higher (*P*_Diet_ =0.04) in FR than during AL (Table 3). Concentrations of ornithine decreased linearly across days (*P*_FR x Day_ < 0.01) while citrulline and BUN (*P*_FR x Day_ < 0.01) increased, all manifesting quadratic increases while their concentration during AL diet remained constant (Table 4). Concentrations of ornithine, citrulline, and NH_3_ were lower after feeding, whereas BUN was higher after feeding regardless of the diet consumed (data not shown).

**Table 3.** Least squares means of the concentrations (μmol/dL) of blood metabolites related with urea cycle and protein tissue mobilization of mid-lactation dairy cows during 5 days on the ad libitum (AL) and feed-restricted (FR) diets.

| Item<sup>1</sup> | Diet |  | SEM | *P* – values<sup>2</sup> |  |  |  |  |
| --- | --- | --- | --- | --- | --- | --- | --- | --- |
|  | AL | FR |  | Diet<sup>3</sup> | Day | Trt x Day | Hour | Trt x Hour |
| 1-MH | 1.02 | 0.97 | 0.09 | 0.99 | <0.01 | 0.3 | 0.01 | 0.89 |
| 3-MH | 0.19 | 0.24 | 0.07 | 0.87 | <0.01 | <0.01 | 0.01 | 0.99 |
| Blood urea N | 426 | 530 | 60 | 0.94 | <0.01 | <0.01 | <0.01 | 0.99 |
| Citrulline | 8.98 | 10.89 | 1.07 | 0.39 | <0.01 | 0.01 | <0.01 | 0.99 |
| NH<sub>3</sub> | 9.76 | 10.49 | 0.74 | 0.04 | 0.11 | 0.16 | <0.01 | 0.52 |
| Ornithine | 4.48 | 4.26 | 0.43 | 0.68 | <0.01 | 0.02 | <0.01 | 0.86 |

<sup>1</sup> α-AAA = α-amino adipic acid; 3-MH = 3-methylhistidine; 1-MH = 1-methylhistidine; PEA = Phosphoethanolamine; EA = Ethanolamine; α-ABA = α-amino butyric acid.
<sup>2</sup>*P*-value of the interactions (Treatment by Day and by Hour) belong to the comparison between the seven treatments.
<sup>3</sup>*P*-value of the orthogonal contrast between the ad libitum (AL) and feed restricted (FR) diets.

**Table 4.**
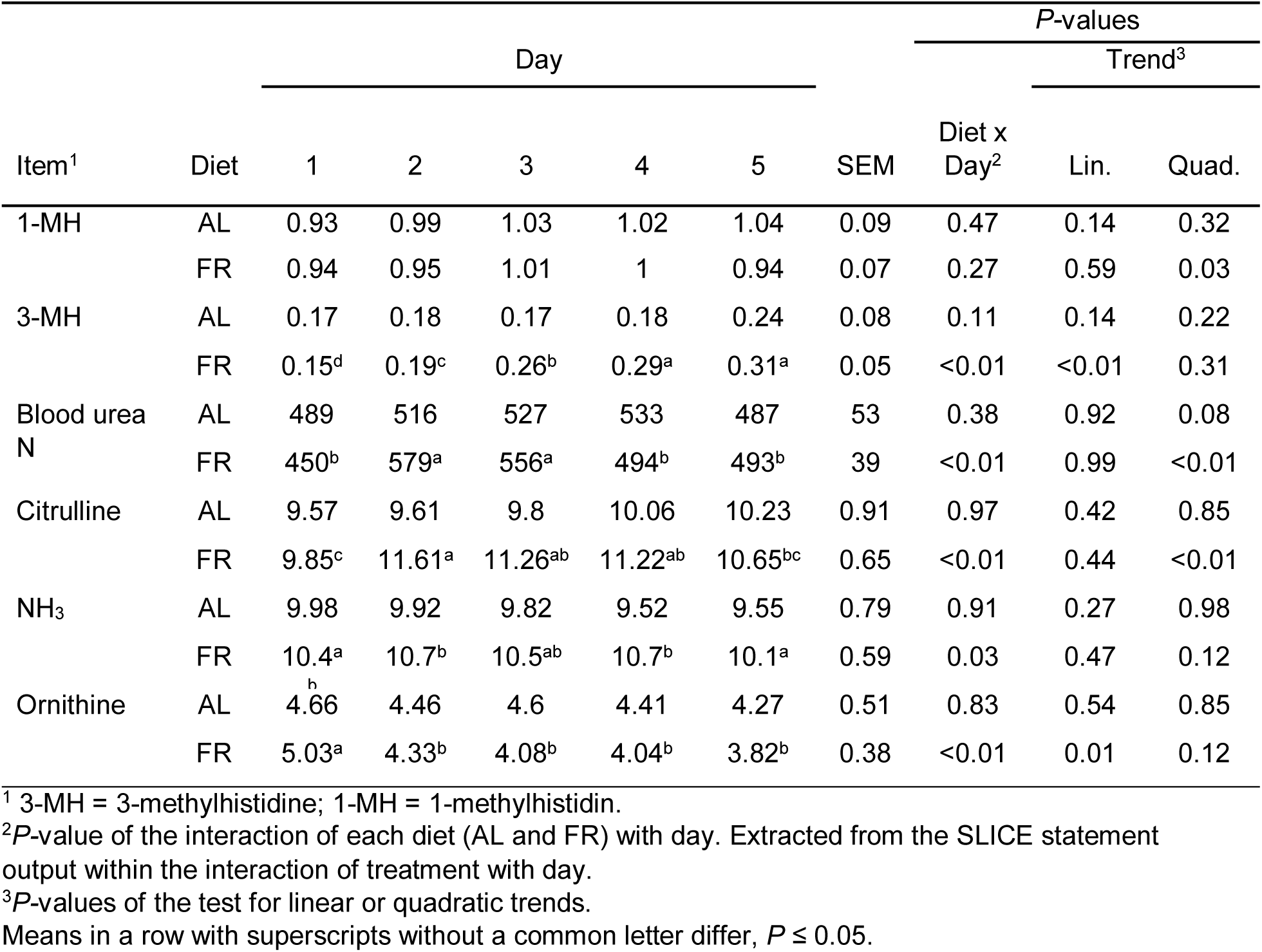
Least squares means of the concentrations (μmol/dL) of blood metabolites related with urea cycle and protein tissue mobilization of mid-lactation dairy cows during 5 days on the ad libitum (AL) and feed-restricted (FR) diets per day.

Other AA such as α-amino adipic acid (**α-AAA**; *P*_Diet_ = 0.11) and amines such as ethanolamine (*P*_Diet_ = 0.12) tended to have greater concentrations during FR (Supplementary Table S1). There was an interaction of FR with day on the daily concentrations of 3-methylhistidine (**3-MH**; *P*_FR x Day_ < 0.01), sarcosine (*P*_FR x Day_ < 0.01), phosphoethanolamine (**PEA**; *P*_FR x Day_ = 0.08), and α-amino butyric acid (**α-ABA**; *P*_FR x Day_ < 0.01), all showing increasing linear trends as the period advanced (Supplementary Table S2). Among these components, we found a significant hour effect for 1-(1-MH) and 3-MH (*P*_Hour_ = 0.01), carnosine (*P*_Hour_ < 0.01), cystathionine (*P*_Hour_ = 0.02), taurine (*P*_Hour_ < 0.01), and α-ABA (*P*_Hour_ = 0.02), all of which had lower concentrations after feeding, and PEA (*P*_Hour_ = 0.05) that was higher relative to feeding time (data not shown).

### Exploratory factor analysis

The principal variables (n = 25) associated with protein metabolism (EAA and NEAA, BUN, NH_3_, and 3-MH as indicator of protein turnover) were reduced to 5 factors that explained 82.2 % of the variance in our data set (Table 5). Factor 1 is a measure of all the EAA except Lys and Arg, plus Ser, Ala, Tyr, Asn, and Pro. Lysine and Arg were correlated with Factor 2 while BUN, 3-MH, and Cit were correlated with Factor 3. Factor 4 was correlated positively with Cys and negatively with NH_3_, Glu, and Asp. Lastly, factor 5 was driven mainly by Gln, Orn, and Gly.

**Table 5.** Loadings in the first five factors extracted by factor analysis of all AA, urea cycle components and 3-methylhystidine in plasma.

| Item | Rotated Factor Loadings<sup>1</sup> |  |  |  |  |
| --- | --- | --- | --- | --- | --- |
|  | Factor | Factor | Factor | Factor | Factor |
| 3-MH | - | - | **0.62** | 0.56 | - |
| Ala | **0.88** | -0.09 | - | -0.23 | - |
| Arg | 0.27 | **0.78** | - | - | 0.4 |
| Asn | **0.79** | 0.28 | -0.26 | - | 0.35 |
| Asp | - | 0.45 | -0.49 | **-0.52** | -0.27 |
| Blood urea N | - | - | **0.83** | - | - |
| Cit | 0.34 | 0.42 | **0.55** | 0.4 | 0.25 |
| Cys | - | 0.3 | - | **0.77** | - |
| Gln | - | - | - | - | **0.85** |
| Glu | - | 0.32 | -0.29 | **-0.7** | - |
| Gly | - | 0.38 | -0.55 | 0.22 | **0.56** |
| Hist | **0.71** | - | - | - | 0.57 |
| Ile | **0.7** | 0.51 | 0.29 | 0.25 | - |
| Leu | **0.86** | 0.25 | - | 0.29 | - |
| Lys | 0.4 | **0.8** | - | - | - |
| Met | **0.71** | 0.48 | - | - | - |
| NH<sub>3</sub> | 0.36 | - | - | **-0.67** | - |
| Orn | 0.53 | 0.46 | - | - | **0.6** |
| Phe | **0.66** | 0.25 | -0.55 | - | - |
| Pro | **0.89** | - | 0.01 | - | 0.33 |
| Ser | **0.63** | - | -0.52 | - | 0.32 |
| Thr | **0.87** | 0.23 | - | - | 0.34 |
| Trp | **0.83** | - | 0.3 | - | - |
| Tyr | **0.89** | - | -0.21 | - | - |
| Val | **0.8** | 0.27 | 0.25 | 0.38 | - |
| Eigenvalue | 10.74 | 4.35 | 2.46 | 1.8 | 1.18 |
| Variance<sup>2</sup>, % | 42.9 | 17.4 | 9.8 | 7.2 | 4.7 |

<sup>1</sup>Correlations between the variable and the factor.
<sup>2</sup>Non-cumulative variance explained by each factor.
Factor loadings smaller than |0.2| were omitted. Factor loadings in bold represent the greatest loading greater than |0.5| within each variable.

## DISCUSSION

The noticeable effect of FR on most AA concentrations reflected their demand during short periods of NNB; therefore, identifying certain AA as potential biomarkers of nutritional status and as targets for supplementation. In this section we discuss those AA that were affected by the main effect of the diet especially the EAA, but it is important to note that whether the diet alone or by through its interaction with day, all AA were affected by feed restriction. The rapid increase of BUN and the balance between the rise in BUN and the decrease in AA in circulation suggest that mobilization of protein tissue may be a quick and important resource to supply energy precursors to maintain milk production and requirements for maintenance.

Since Pro is one of the AA with greatest abundance in the body, due to its function in collagen synthesis, its demand may overcome supply in certain scenarios. In a nutrient stress situation, Pro can be metabolized to yield Glu and then α-ketoglutarate (**α-KG**), which can participate in gluconeogenesis. The Pro concentration in plasma usually declines around parturition and then increases quickly in early lactation (Maeda et al., 2012). However, Laeger et al. (2012) and Kuhla et al. (2011) did not find any variation of Pro after feed restriction.

Enzymatic conversion from Phe to Tyr is increased by glucagon and inhibited by insulin, while catabolism of Tyr is induced by cortisol and other glucocorticoid hormones. Therefore, during FR it seems that a greater degradation of Phe and Tyr may have be triggered by the decrease in insulin (Ansia et al. unpublished results) and the likely increase in plasma cortisol concentration (Toerien and Cant, 2007), driving their catabolism towards gluconeogenesis from these AA and reducing their protein synthesis potential (Doepel et al., 2016). Laeger et al. (2012) found a tendency for a lower plasma concentration of Tyr and a lower Tyr concentration in cerebrospinal fluid, which would suggest a potential central anorexic effect. Tyrosine is also a precursor of thyroid hormones in the thyroid gland, as well as neurotransmitters (catecholamines) and hormone precursor in neurons and adrenal medulla, respectively. Among catecholamines, adrenaline, noradrenaline, and dopamine have a recognized anorexic effect that would help counteract hunger. Although during FR periods synthesis of thyroid hormones seems to decrease, catecholamines in circulation increase to reduce milk production and increase lipogenesis (Salin et al., 2017).

Laeger et al. (2012) reported a similar reduction (−12%) to ours in plasma Thr concentration and an even larger decrease in the cerebrospinal fluid concentration of Thr (−24%), which suggests that Thr could also act as a signal for feed intake regulation. Alike most of the EAA, Thr concentration also decreases as calving approaches and increases as lactation advances (Larsen et al., 2015). Reduction of Thr has been reported consistently in short periods of FR (Baird et al., 1972; Kuhla et al., 2011). Addition of extra Thr through abomasal infusions in early lactation did not have a positive effect, but neither did its deletion cause a negative consequence (Doepel et al., 2016). This outcome suggests that even on a low protein intake, the cow would be able to meet its Thr requirements.

An increase or no variation in Gly concentration during NNB has been reported in the literature during short-term feed restriction periods (Kuhla et al., 2011; Laeger et al., 2012) or during the transition period (Larsen and Kristensen, 2009), whereas Baird et al. (1972) reported a non-significant decrease (−27%) after 6 d of complete starvation. This common observation could be due to a reduced mammary (Raggio et al., 2006) and hepatic (Larsen and Kristensen, 2009) uptake, and to a net release triggered by tissue mobilization. Glycine is the backbone of the basic unit that form connective tissue, and is also one of the AA with greatest abundance in muscle in its free form (Meijer et al., 1995). Sarcosine is a direct precursor of Gly and its concentration was also increased during FR in this study; therefore, it is likely that the sarcosine pathway was active during FR.

Lysine uptake is greater than its milk output and occurs mostly in the mammary gland (Mepham, 1982). Oxidation of Lys provides the N-group for synthesis of other NEAA such as Arg and Pro. An increase in Lys concentration during feed restriction has been commonly reported (Baird et al., 1972; Kuhla et al., 2011; Laeger et al., 2012). The decrease of milk protein production suggests that Lys would not be demanded for this anabolic process since the mammary gland has the capacity to alter AA uptake to meet its requirements (Mackle et al., 2000). Guo et al. (2017) found that mammary Lys clearance and the ratio mammary uptake:milk output only decreased significantly when all Lys supply was removed. This result indicates that there could be a compensatory mechanism to supply a certain amount of Lys to the mammary gland even with low nutritional supply. In fact, post-hepatic supply is greater than mammary uptake regardless of the portal net absorption of Lys, and mammary uptake decreases by only a small proportion relative to a reduced post-hepatic supply (Lapierre et al., 2009). A reduction in hepatic uptake could explain why plasma Lys concentration tended to be higher during FR, if the mammary uptake was still maintained. However, uptake of Lys for milk protein synthesis could have been reduced in favor of catabolism toward synthesis of other NEAA such as Arg and Pro (Lapierre et al., 2009; Guo et al., 2017). The most common pathway for Lys catabolism is the saccharopine pathway, which uses α-amino adipic acid (α-AAA) as intermediate. High concentrations of α-AAA despite the elevated concentrations of Lys during FR could in fact suggest that hepatic and extrahepatic oxidation are part of the regulatory mechanism to control the excess of Lys in circulation (Tucker et al., 2017).

Mammary gland takes up Arg at a higher rate than milk output as well. Its catabolism contributes to milk protein synthesis and as AA precursor (Clark et al., 1978). As demonstrated in vitro, Arg plays a crucial role in protein synthesis as regulator of transcription in casein and m-TOR pathway genes in mammary epithelial cells (Wang et al., 2014). If regulation of the mammary extraction efficiency involves specific AA transporters (Mackle et al., 2000), Lys and Arg could have been equally affected during FR because they share almost exclusively (with ornithine to some extent) the same unique cationic transporter (Y^+^) across mammary tissue membranes (Baumrucker, 1984), which would explain the greater plasma concentration of both AA during FR. Therefore, the reduction in milk protein synthesis caused by the endocrine changes during NNB could be either triggered by or augmented by disabled transport of Lys and Arg into the mammary gland. In fact, Rius et al. (2010) found that only abomasal infusion of starch (with that concomitant raise of insulin and IGF-1 concentrations) rather than casein during feed restriction was able to stimulate mammary AA uptake and milk protein synthesis despite excess availability of AA.

In other studies similar to ours, His concentration did not decrease during feed restriction (Baird et al., 1972; Laeger et al., 2012), or even was higher (Kuhla et al., 2011). However, in early lactation His seems to undergo a high uptake by the mammary gland (Larsen et al., 2015), causing a prolonged deficiency since arterial levels did not return to pre-calving concentrations during the time-frame of most studies (Meijer et al., 1995; Doepel et al., 2002). Histidine in body tissue is methylated when the muscle contractile proteins are broken down during protein mobilization. This new form, 3-MH, cannot be recycled for protein synthesis and thus has to be excreted, becoming an optimal index of tissue protein turnover (Doepel et al., 2002). However, 3-MH plasma concentration is not a proportional indicator of amount of muscle mobilized because its incomplete recovery in urine suggests a pool of non-protein bound 3-MH in muscle (Lobley, 1998). 1-Methylhistidine is another methylated form of His that in combination with β-alanine forms anserine, which is a methylated form of carnosine present in skeletal muscle that also has anorexigenic effects (Tomonaga et al., 2004). While 3-MH and 1-MH must be excreted, β-alanine can be catabolized to yield acetyl-CoA in tissues. During the FR diet, daily averages of carnosine (−9%) and β-alanine (−15%) showed similar patterns of decrease that matched the increase of 1-MH (+8%). The same opposite pattern across time between carnosine and 1-MH was reported by Laeger et al. (2012). These results could indicate that carnosine in muscle likely was degraded during FR, releasing β-alanine that also would be catabolized and 1-MH that would be excreted.

Since dietary N intake was reduced, the increase of BUN during FR indicates that the urea cycle was in motion to eliminate excess N produced by AA catabolism or as result of N recycling from the digestive tract to use endogenous urea as N source. Higher BUN concentrations also were reported during feed-restriction conditions in other experiments (Loor et al., 2007; Kuhla et al., 2011). The rapid response of BUN to FR probably could have contributed to hide this effect in other studies with FR (Carlson et al., 2006; Laeger et al., 2012), or even during the onset of lactation (Zhu et al., 2000), where sampling was as frequent. In addition, plasma ammonia concentration is probably a better indicator of AA catabolism than BUN during the transition period because expression of some of the enzymes involved in ureagenesis is decreased at calving and does not seem to react accordingly with the requirements at the onset of lactation (Hartwell et al., 2001).

Both Glu and Asp play an important role as they are involved in the urea and tricarboxylic acid cycles, both highly active because of their implications for AA catabolism and gluconeogenesis during periods of NNB (Noro and Wittwer, 2012). Glutamate provides α-KG, an intermediate in the tricarboxylic acid cycle, and the α-amino N required to synthetize Asp from oxaloacetate. The synchronized and opposite patterns across days for BUN and Gln during FR seems to elucidate that urea formation was the predominant method at the beginning of dietary restriction, probably due to the greater detoxification capacity of ammonia at lower energetic cost (Reynolds, 2006). However, as days pass urea synthesis seemed to decrease and Gln synthesis increased, suggesting a switch towards the low-capacity, high-affinity method to detoxify ammonia as Gln (Lobley et al., 1995). This shift could have been established to save bicarbonate ions and maintain an adequate cation-anion balance (Meijer et al., 1993).

The increase in BUN coincided with the steepest decrease in plasma concentrations of most AA. We acknowledge that these estimates are not quantitative because fluxes of compounds and differential half-lives in circulation were not measured. However, accounting for the balance of N compounds measured in plasma by comparing incremental area under the curve (IAUC) of the N-elements whose concentration increased from d 1 to d 5 (N-AA^+^) and BUN, with those whose concentration decreased (N-AA^-^), there seemed to be a shortfall of amino-N sources for the increase in BUN. Considering the limitations of this rough comparative method, during AL we observed a quite well balanced equilibrium between the positive IAUC of N-AA^+^ (56 μg·d) and BUN (735 μg·d) in plasma, and the negative IAUC of N-AA^-^ (−544 μg·d). However, during FR the IAUC of BUN (2,266 μg·d) and N-AA^+^ (207 μg·d) increased when compared with the AL diet to a much greater degree (2x) than the IAUC of N-AA^-^ (−1,135 μg·d).

During short periods of feed restriction hepatic uptake of lactate seems to not play a great role as a precursor for gluconeogenesis (Laeger et al., 2012), unlike in early lactation (Larsen and Kristensen, 2012). Therefore, tissue mobilization and peptides in circulation probably could have contributed to the AA turnover and catabolism, being then the source of that “extra” BUN during FR. In accordance with the literature, plasma proteins, liver protein, and gastrointestinal tract tissue would be the first mobilized (Swick and Benevenga, 1977), while skeletal muscle breakdown probably would be more intense at the end of the period as the increasing concentrations of 3-MH and α-AAA in plasma indicated in our study.

### Exploratory factor analysis

Factor analysis allowed us to reduce some of the principal variables involved in protein metabolism (individual AA, NH_3_, BUN, and 3-MH) into a reduced number of factors identifying underlying correlations among those variables. Loadings of factor 1 confirm the common effect of FR over all the EAA except Lys and Arg, plus the NEAA Ser, Ala, Tyr, Asn, and Pro, whose concentrations were reduced during FR. Factor 2 loadings demonstrate that FR provoked a different effect over Lys and Arg than over the majority of EAA, and emphasizes their mutual correlation as previously discussed. Due to the required inter-organ coordination during ureagenesis, Cit concentration, rather than Arg, probably determines better the concentration of BUN since both were grouped in factor 3. In addition, the correlation of BUN with 3-MH within factor 3 reveals that the catabolism of tissue could be an important source of the N captured and excreted as urea. Factor 4 indicates the different evolution of the increasing concentrations of Cys, and elucidates the coordinated demand for Asp and Glu in the urea cycle as shown by their correlation with NH_3_ (Noro and Wittwer, 2012). Lastly, grouping of Gln and Gly within factor 5 could suggest a similar effect over their release from muscle upon tissue breakdown. Glutamine has the highest concentration in muscle as free AA (Meijer et al., 1995) and Gly is omnipresent in the tri-peptide that forms the basic unit in connective tissue; therefore, their higher concentration could be indicative of a release upon muscle degradation (Doepel et al., 2002).

## CONCLUSIONS

The decrease in insulin concentration during FR coordinately triggered lipid and AA mobilization and oxidation as reflected by the greater NEFA and BUN concentrations in plasma and milk. The rapid increase in BUN was paired with a decrease in concentrations of all EAA except Lys and Arg, plus the NEAA Ser, Ala, Tyr, Asn and Pro, which were the AA that better represented the effect of FR in plasma. In contrast, the increase in Lys and Arg concentrations could be a response to the reduced requirements for milk protein synthesis. Protein tissue mobilization and modulation of tissue insulin sensitivity may be important mechanisms to supply N and energy substrates for metabolic fuel, and NEFA and glucose for milk synthesis, respectively, during a short period of restricted DMI. Results from our model highlight 1) the need for more research to better understand the role of protein tissue catabolism and endocrine regulation, and 2) that supplementation of certain AA could improve or be a proxy of health and performance around the transition period or even during short periods of restricted feed intake.

## Acknowledgments

The authors thank Ajinomoto Co., Inc. Tokyo, Japan for providing the funds for this experiment.

## Ethics statement

All procedures involving animals were approved by the Institutional Animal Care and Use Committee at the University of Illinois at Urbana-Champaign (protocol #15167).

