## Supplemental file for "Feed restriction in mid-lactation dairy cows. II: Effects on protein metabolism-related blood metabolites"

### Supplemental tables

**Supplementary Table S1**. Concentrations (μmol/dL) of blood metabolites related with N metabolism of mid-lactation dairy cows during 5 days on the ad libitum (AL) and feed-restricted (FR) diets per day.

| | Diet | | | *P* – values[2] | | | | |
|---|---|---|---|---|---|---|---|---|
| Item[1] | AL | FR | SEM | Diet[3] | Day | Trt x Day | Hour | Trt x Hour |
| Carnosine | 1.29 | 1.09 | 0.19 | 0.37 | <0.01 | 0.01 | <0.01 | 0.99 |
| Cystathionine | 0.14 | 0.15 | 0.03 | 0.68 | <0.01 | <0.01 | 0.02 | 0.88 |
| Ethanolamine | 0.40 | 0.37 | 0.04 | 0.12 | 0.02 | 0.03 | 0.29 | 0.97 |
| Hydroxyproline | 1.03 | 1.13 | 0.20 | 0.37 | 0.09 | 0.89 | <0.01 | 0.5 |
| PEA | 2.02 | 2.15 | 0.15 | 0.51 | 0.02 | <0.01 | 0.05 | 0.97 |
| Phosphoserine | 1.42 | 1.33 | 0.09 | 0.48 | <0.01 | <0.01 | 0.06 | 0.99 |
| Sarcosine | 0.56 | 0.51 | 0.14 | 0.8 | <0.01 | <0.01 | 0.2 | 0.45 |
| Taurine | 6.03 | 5.76 | 1.04 | 0.55 | <0.01 | <0.01 | <0.01 | 0.85 |
| α-AAA | 0.70 | 0.92 | 0.10 | 0.11 | <0.01 | 0.27 | 0.53 | <0.01 |
| α-ABA | 1.65 | 1.87 | 0.20 | 0.43 | <0.01 | <0.01 | 0.02 | 0.99 |
| β-Alanine | 0.22 | 0.24 | 0.06 | 0.55 | <0.01 | <0.01 | 0.07 | 0.91 |

[1]PEA = Phosphoethanolamine; α-AAA = α-amino adipic acid; α-ABA = α-amino butyric acid.
[2]*P*-value of the interactions (Treatment by Day and by Hour) belong to the comparison between the seven treatments.
[3]*P*-value of the orthogonal contrast between the ad libitum (AL) and feed restricted (FR) diets.

**Supplementary Table S2.** Concentrations of blood metabolites related with nitrogen metabolism of mid-lactation dairy cows during 5 days on the ad libitum (AL) and feed-restricted (FR) diets per day.

| | | | | | | | | *P*-values | | |
|---|---|---|---|---|---|---|---|---|---|---|
| | | Day | | | | | | | Trend[3] | |
| Item[1] | Diet | 1 | 2 | 3 | 4 | 5 | SEM | Diet x Day[2] | Lin. | Quad. |
| Carnosine | AL | 1.2 | 1.28 | 1.27 | 1.33 | 1.36 | 0.21 | 0.49 | 0.13 | 0.99 |
| | FR | 1.17 | 1.12 | 1.05 | 1.06 | 1.05 | 0.14 | 0.39 | 0.1 | 0.31 |
| Cystathionine | AL | $0.14^{bc}$ | $0.14^{b}$ | $0.16^{a}$ | $0.16^{ab}$ | $0.13^{c}$ | 0.03 | <0.01 | 0.86 | 0.05 |
| | FR | $0.15^{a}$ | $0.15^{ab}$ | $0.15^{ab}$ | $0.15^{ab}$ | $0.14^{b}$ | 0.02 | 0.24 | 0.31 | 0.93 |
| Ethanolamine | AL | 0.38 | 0.38 | 0.41 | 0.42 | 0.41 | 0.04 | 0.73 | 0.27 | 0.64 |
| | FR | $0.37^{ab}$ | $0.36^{b}$ | $0.41^{a}$ | $0.36^{b}$ | $0.36^{b}$ | 0.04 | <0.01 | 0.62 | 0.22 |
| Hydroxyproline | AL | 1.09 | 0.96 | 1.05 | 1.07 | 1.01 | 0.19 | 0.68 | 0.92 | 0.92 |
| | FR | $1.11^{ab}$ | $1.22^{a}$ | $1.15^{ab}$ | $1.23^{a}$ | $1.08^{b}$ | 0.15 | 0.13 | 0.88 | 0.28 |
| PEA | AL | 2.08 | 1.97 | 2.03 | 1.99 | 2.01 | 0.17 | 0.85 | 0.71 | 0.63 |
| | FR | $2.09^{b}$ | $2.05^{b}$ | $2.17^{ab}$ | $2.13^{b}$ | $2.29^{a}$ | 0.13 | 0.08 | 0.04 | 0.31 |
| Phosphoserine | AL | $1.55^{d}$ | $1.47^{cd}$ | $1.42^{bc}$ | $1.35^{ab}$ | $1.30^{a}$ | 0.09 | <0.01 | <0.01 | 0.83 |
| | FR | 1.36 | 1.35 | 1.32 | 1.31 | 1.31 | 0.07 | 0.84 | 0.18 | 0.67 |
| Sarcosine | AL | 0.55 | 0.56 | 0.58 | 0.58 | 0.53 | 0.15 | 0.6 | 0.89 | 0.39 |
| | FR | $0.45^{b}$ | $0.48^{b}$ | $0.52^{ab}$ | $0.54^{a}$ | $0.57^{a}$ | 0.1 | <0.01 | 0.01 | 0.88 |
| Taurine | AL | 6.07 | 6.1 | 5.89 | 6.2 | 5.91 | 1.15 | 0.92 | 0.91 | 0.96 |
| | FR | 5.92 | 5.73 | 5.58 | 5.6 | 5.99 | 0.81 | 0.96 | 1 | 0.38 |
| α- AAA | AL | 0.67 | 0.64 | 0.64 | 0.64 | 0.7 | 0.12 | 0.97 | 0.86 | 0.53 |
| | FR | $0.78^{bc}$ | $0.72^{c}$ | $0.79^{c}$ | $0.97^{ab}$ | $1.02^{a}$ | 0.11 | 0.03 | 0.01 | 0.1 |
| α-ABA | AL | 1.47 | 1.65 | 1.77 | 1.79 | 1.65 | 0.26 | 0.16 | 0.4 | 0.02 |
| | FR | $1.50^{d}$ | $1.71^{c}$ | $1.93^{b}$ | $2.10^{a}$ | $2.14^{ab}$ | 0.18 | <0.01 | <0.01 | 0.1 |
| β-Alanine | AL | 0.23 | 0.21 | 0.23 | 0.22 | 0.22 | 0.06 | 0.85 | 0.78 | 0.92 |
| | FR | 0.25 | 0.23 | 0.22 | 0.26 | 0.26 | 0.05 | 0.39 | 0.42 | 0.14 |

[1] PEA = Phosphoethanolamine; α-AAA = α-amino adipic acid; α-ABA = α-amino butyric acid.

[2]*P*-value of the interaction of each diet (AL and FR) with day. Extracted from the SLICE statement output within the interaction of treatment with day.

[3]*P*-values of the test for linear or quadratic trends.

Means in a row with superscripts without a common letter differ, $P \leq 0.05$.
